# A dataset of simultaneous two-photon calcium imaging and auditory discrimination behavior

**DOI:** 10.64898/2026.04.28.721534

**Authors:** Binjie Hong, Li Li, Yongzhen Yu, Yu Xin, Yuan Zhang, Ninglong Xu, Tielin Zhang

## Abstract

The emerging field of artificial intelligence for neuroscience relies heavily on high-quality, standardized datasets. Data linking high-dimensional neural population dynamics with complex behavior is crucial for advancing the field of brain-computer interfaces. However, few existing datasets provide these paired modalities in a standardized, AI-ready format, limiting their immediate utility for computational modeling. Here, we present a large-scale dataset comprising simultaneous two-photon calcium imaging of the primary auditory cortex (A1) and detailed behavioral records in mice. The animals performed an auditory two-alternative forced-choice (2AFC) categorization task, classifying tones as low or high frequency by licking left or right water ports. The raw neural and behavioral data were preprocessed, synchronized, and formatted into trial-aligned multi-dimensional tensors. Comprehensive validation confirms that the recorded populations exhibit consistent, high-fidelity temporal dynamics and distinct frequency tuning across all experimental conditions. To evaluate the dataset’s capacity for neural decoding, we benchmarked it across diverse machine learning architectures and deep learning networks. By bridging biological recordings and behavior, this open-access dataset serves as a valuable benchmark for developing novel decoding algorithms and testing brain-inspired computational models.

## Background & Summary

Recent advances in two-photon calcium imaging have revolutionized neuroscience by enabling the simultaneous recording of large neuronal populations with single-cell resolution[1, 2, 3, 4]. This optical recording approach provides a unique window to link high-dimensional neural dynamics with complex behavioral outputs. As the primary auditory cortex (A1) has been viewed as a sensory hub for feature extraction, recent research demonstrates its deep involvement in non-sensory cognitive processes[5, 6]. Contemporary studies reveal that A1 encodes variables extending far beyond sound frequency, including reward prediction [7], motor planning [8], and decision-making variables [9]. The mouse, with its robust auditory learning capabilities and accessible genetic toolkit for cell-type specific targeting, serves as an ideal model organism to dissect these circuit mechanisms[10, 11].

While modern neuroscience generates large-scale biological data, the lack of precise alignment with behavior limits its immediate utility for NeuroAI, which bridges neuroscience and artificial intelligence [12]. Specifically, high-quality, standardized datasets pairing decision-making behaviors with population recordings remain scarce. Since most existing datasets require extensive preprocessing, the application of advanced neural decoding models is severely hindered. To address this gap, we present a large-scale aligned dataset of mouse A1 activity during auditory discrimination tasks.

The dataset is designed to investigate auditory processing mechanisms across different acoustic frequency contexts. The recordings span three distinct auditory frequency bands to ensure generalizability: a standard subset (8–32 kHz) for benchmarking auditory discrimination, alongside low-shift (4–16 kHz) and high-shift (7–28 kHz) conditions to investigate processing across different spectral ranges.

Rigorous technical validation confirmed the high biological fidelity of the dataset at both the population and single-neuron levels. We capture the frequency selectivity of individual neurons and the characteristic tonotopic organization. Since all neural and behavioral data were preprocessed, synchronized, and formatted into trial-aligned tensors, we also conduct a comparative analysis of neural decoding. Both traditional machine learning and deep learning models are assessed, including Support Vector Machines (SVM), Multilayer Perceptrons (MLP), Convolutional Neural Networks (CNN), and Long Short-Term Memory (LSTM) networks. They decode population dynamics and explore the neural computations underlying sensory-guided decision-making[13, 14].

## Methods

### Experiment Preparation

All experimental procedures were conducted in strict compliance with protocols approved by the Animal Care and Use Committee of the Institute of Neuroscience, Chinese Academy of Sciences. Data were primarily acquired from male C57BL/6J mice (Shanghai Laboratory Animal Center, SLAC). Animals were 9–10 weeks old at the onset of behavioral training and 10–14 weeks old during the two-photon calcium imaging sessions.

To motivate behavioral performance, mice were placed on a controlled water restriction schedule. On non-training days, mice received a daily ration of 1 mL of water. On training days, mice participated in experimental sessions lasting 1–2 hours, during which they obtained approximately 0.5–1.0 mL of water as behavioral rewards. The body weight of each animal was monitored daily to ensure it was maintained above 80% of their pre-restriction baseline weight. For a comprehensive description of the surgical procedures and viral injection coordinates, please refer to the companion article [15].

### Surgery

Mice were anesthetized with isoflurane (1–2%) during all surgical procedures. A craniotomy (approx. 2 mm in diameter) was performed over the left primary auditory cortex (A1), with the dura mater left intact. To label cortical neurons, the calcium indicator AAV-hSyn-GCaMP6s was microinjected (50–150 nL per site; 3–4 sites per animal) using a pulled glass micropipette (25–30 µm O.D.) controlled by a hydraulic manipulator (MO-10, Narashige) and positioned via a Sutter MP-225 manipulator. Following viral injection, a custom double-layered cranial window was implanted to seal the craniotomy. This optical window consisted of a 200-µm-thick glass coverslip (inner diameter: 2 mm) bonded to a larger coverslip (outer diameter: 5 mm) using UV-curable adhesive. The window was secured to the skull using dental cement (Jet Repair Acrylic, Lang Dental Manufacturing). A titanium head-post was affixed to the skull to facilitate head fixation during two-photon imaging. Post-surgery, mice were allowed to recover for at least 7 days before the commencement of water restriction and behavioral training.

### Behavioral Task

Experiments were performed in custom double-walled sound-attenuating chambers. Head-fixed mice were restrained in acrylic tubes and provided with bilateral capacitive-sensing lick ports for water reward[16]. The behavioral task was controlled by a custom real-time system (“PX-Behavior”) based on an Arduino microcontroller, which managed stimulus delivery and logged events via Python software. Crucially, behavioral trials were synchronized with two-photon image acquisition using digital TTL pulses. Auditory stimuli were delivered via an electrostatic speaker (ES1, Tucker-Davis Technologies) positioned to the right of the animal. The acoustic system was calibrated using a free-field microphone to ensure a flat spectrum (±5 dB) across 3–60 kHz, with 5 ms cosine ramps applied to all tone onsets and offsets to minimize spectral splatter.

Mice were trained on an auditory-guided two-alternative forced-choice (2AFC) categorization task under water restriction [17]. The experimental workflow and trial structure are illustrated in Figure 1. Each trial began with a random delay (0.5–1.0 s) to prevent temporal prediction, followed by the presentation of a pure tone stimulus (duration: 300 ms; intensity: 70–75 dB SPL). A 500 ms post-stimulus delay was imposed before the 3 s response window, during which mice reported their decision. As shown in Figure 1, licking the left port for low-frequency tones or the right port for high-frequency tones triggered a water reward (6 µL), whereas incorrect choices resulted in a 2–6 s time-out punishment. Noticeably, data collection commenced once mice achieved stable performance with 75-85% accuracy.

**Figure 1:**
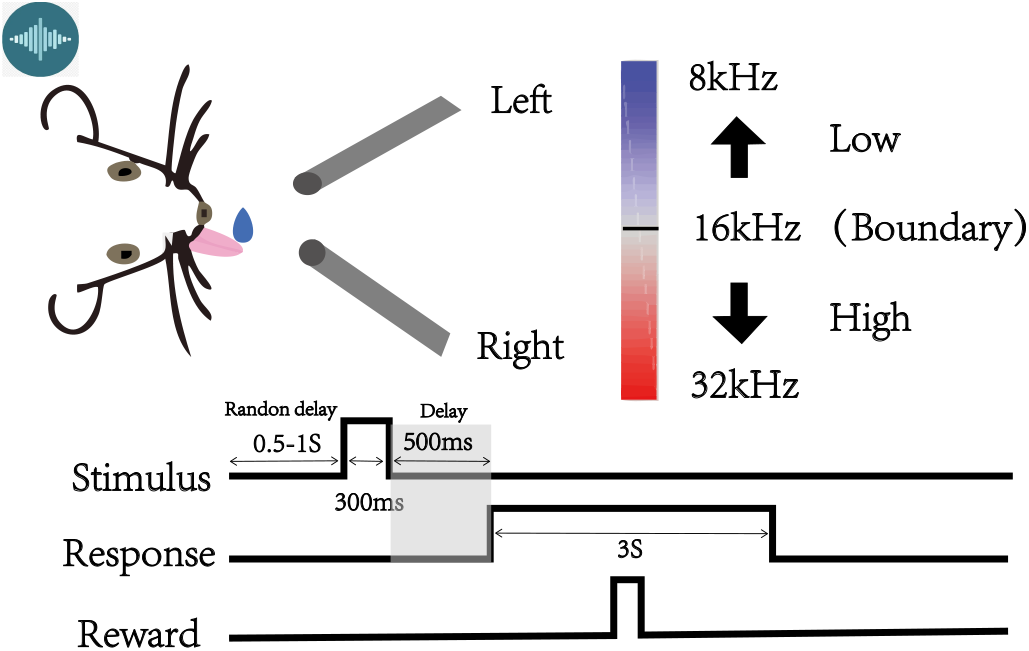
Mice were head-fixed and trained to classify tone frequencies into low or high categories by licking the left or right water port. Each trial followed a strict timeline: a random delay, sound stimulus, a mandatory delay, and a response window during which correct choices triggered a water reward.

### Data Acquisition

To bridge the gap between raw biological recordings and computational modeling, we developed a standardized data processing pipeline as illustrated in Figure 2 that integrates two parallel streams: neural tensor transformation and behavioral vectorization. Ultimately, the raw recordings are transformed into paired datasets, fusing standardized neural tensors shaped as *N*_trials_ *× N*_cell_ *× N*_frames_ with corresponding behavioral labels.

**Figure 2:**
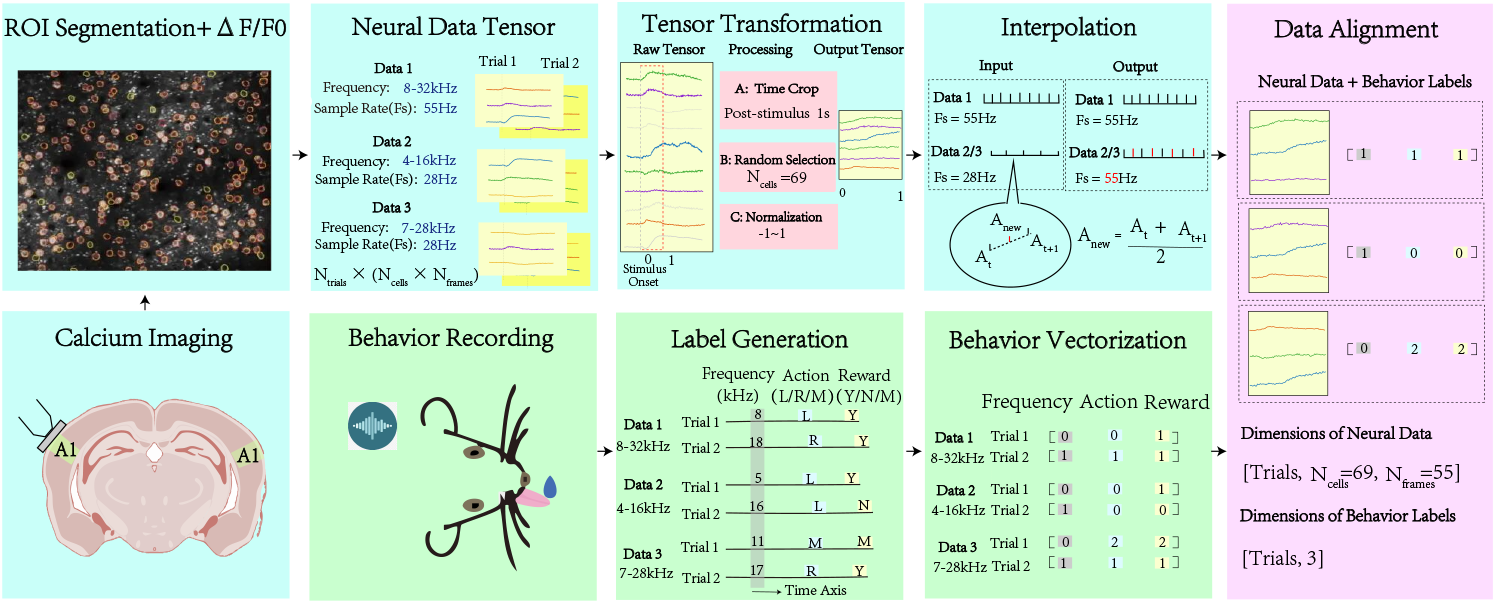
The pipeline aligned raw biological signals and corresponding behaviors. Calcium imaging stacks are segmented to extract fluorescence traces. These neural tensors are then standardized via time-cropping, spatial subsampling to unify feature dimensions, normalization, and temporal interpolation (blue panels). Simultaneously, raw behavioral logs are parsed and vectorized into numerical labels representing frequency, actions, and rewards (green panels). Finally, the processed neural tensors and behavioral vectors are fused to create paired datasets for downstream applications (purple panel).

#### Neural Tensor Transformation and Standardization

The two-photon calcium imaging data, including relative fluorescence changes and deconvolved spike probabilities, were obtained following standard ROI segmentation and preprocessing procedures as detailed in the companion study [15]. These extracted neural traces were temporally synchronized with behavioral events and initially organized into 3D tensors in the shape of *N*_trials_ *×N*_cells_*× N*_frames_.

To ensure compatibility across subsets on different conditions and prepare the data for down-stream machine learning applications, we implemented a sequential standardization procedure.

First, the neural traces were time-cropped to a fixed 1-second window immediately following the stimulus onset, which isolates the critical decision-making epoch. Next, we aim to align the spatial feature dimensions across recording sessions with variable cell yields, hence we randomly subsampled the neural population to a fixed size of 69 neurons per trial. The number of neurons is determined by the minimum yield observed across all experiments. Furthermore, to resolve discrepancies in data acquisition speeds, traces originally sampled at 28 Hz in the shifted-frequency groups were temporally upsampled to match the 55 Hz rate of the standard condition via linear interpolation, expressed as:

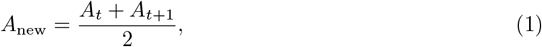

where *A* denotes the neural activity value (Δ*F/F*_0_). *A*_*t*_ and *A*_*t*+1_ represent the recorded signal amplitudes at consecutive frames in the 28 Hz time series, and *A*_new_ is the interpolated midpoint inserted to approximate the 55 Hz temporal resolution. Finally, we normalized neural activity and bound the values within a consistent range [*−*1, 1] for modeling stability. The standardized tensors underwent *Z*-score scaling followed by a hyperbolic tangent (tanh) transformation:

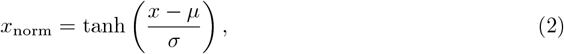

where *x* is the processed Δ*F/F*_0_, and *µ* and *σ* are the mean and standard deviation of the pre-stimulus baseline period, respectively.

### Behavioral Vectorization

To facilitate computational tasks, raw behavioral records were parsed and encoded into structured numerical vectors. For each experimental trial indexed as *i*, the 2AFC task label was mapped to a vector *B*_*i*_ = [*Frequency, Action, Reward*]. As detailed in Table 1, the stimulus category was binarized as 0 and 1 based on the specific frequency boundary of each experimental condition. The mouse’s action choice was encoded as 0 for licking left, 1 for licking right, and 2 for missing. The trial outcome of reward was evaluated as 1 for correctness, 0 for error, and 2 for missing actions. This standardized formatting provides a clear, trial-resolved ground truth across all experimental groups.

**Table 1:** Metadata description for stimulus and behavioral variables.

| Field Name | Description | Value Encoding |
| --- | --- | --- |
| *Frequency* | Auditory stimulus classification based on task boundary. Data consists of three groups with logarithmically spaced frequencies:<br>Standard: 8–32 kHz range (Boundary: 16 kHz)<br>Low-Shift: 4–16 kHz range (Boundary: 8 kHz)<br>High-Shift: 7–28 kHz range (Boundary: 14 kHz) | 0: Low Frequency (< Boundary)<br>1: High Frequency (> Boundary) |
| *Action* | Mouse licking response recorded during the decision window. | 0: Left Lick<br>1: Right Lick<br>2: Miss (No Response) |
| *Reward* | Evaluation of trial success, defining the correctness as (*Frequency* =0 & *Action*=0) or (*Frequency* =1 & *Action*=1). | 0: Error (No Reward)<br>1: Correct (Reward)<br>2: Miss (No Response) |

## Data Records

To facilitate cross-platform investigation, we provide the data in a processed, structured format. The dataset is organized by experimental sessions, and files are provided in MATLAB (.mat) format to maximize accessibility. The complete dataset is publicly available through the Chinese Academy of Sciences Data Center (https://www.braindatacenter.cn).

### File Structure

The raw neural data is stored in the main directory, organized by experimental groups: 8 32kHz Data/, 4 16kHz Data/, and 7 28kHz Data/. Each group contains multiple session files named as Sess1 data.mat. Each file includes two tensors: Frequency Action Reward of shape [Trials, Cells, Frames] and Labels of shape [Trials, Labels]. The Frequency Action Reward tensor provides the neural activity for model input, while the Labels tensor supplies the behavioral labels for training.

Figure 3 summarizes the quantitative characteristics of the three experimental groups. The 8-32 kHz group comprises 21 sessions, with averages of 220 trials, 124 cells, and 344 frames per session. In comparison, the 4–16 kHz and 7–28 kHz groups each contained 12 sessions, exhibiting similar scales in average trials, cells, and frames.

**Figure 3:**
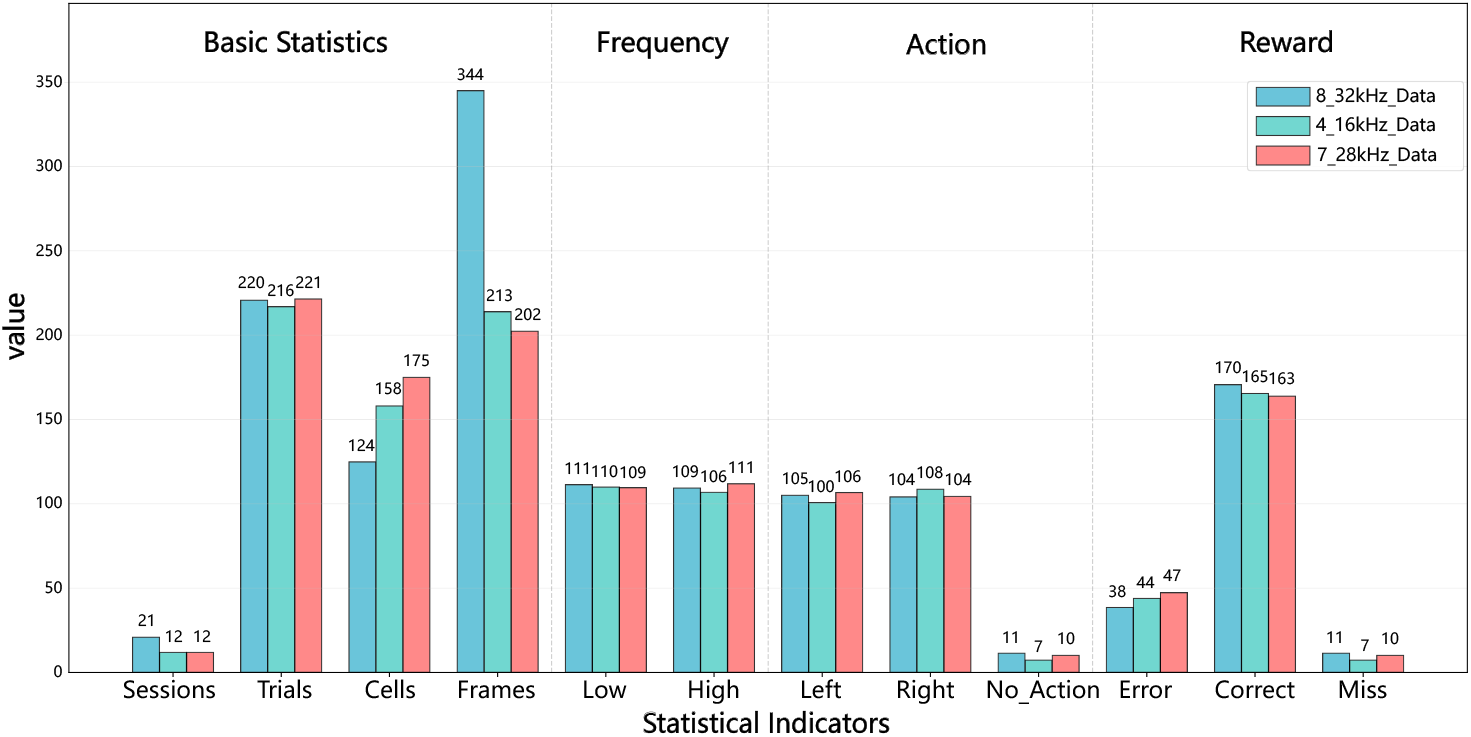
Statistics across the three experimental groups. The bar chart displays the session count and the average per-session values for three core dataset statistics, alongside the average per-session distribution of the three prediction targets: low or high frequency, actions of licking left or right, and reward.

The label distributions are consistent across groups. The tone frequency labels have balanced counts of low and high tones. Action labels are balanced per session, with minimal no-action trials. Correct reward trials are the most frequent, followed by incorrect licking and missing rewards. This balanced distribution supports merging all three groups for subsequent modeling.

## Technical Validation

### Data Consistency

#### Neural Response in Diverse Frequency Conditions

To validate how the brain reacts in different-frequency stimuli, we analyzed the population dynamics of sound-evoked activity across all three experimental groups (Figure 4). To isolate neurons with robust and reliable responses, we applied a two-step statistical filter comparing a 1.5 s post-stimulus window against a 0.5 s pre-stimulus baseline. First, a paired t-test (*P <* 0.05) was utilized to verify that the evoked activity was statistically significant rather than a random baseline fluctuation. Second, a peak Z-score threshold (|*Z*| *>* 0.5) was applied to ensure a sufficiently high signal-to-noise ratio, thereby excluding neurons with statistically significant but biologically negligible response magnitudes. Consequently, neurons exhibiting excitation with a peak *Z*-score greater than 0.5 were classified as activated neurons, whereas those with inhibition and a peak *Z*-score less than −0.5 were classified as the suppressed. The temporal dynamics of these populations were then visualized using grand-averaged, *Z*-scored fluorescence traces, while their relative proportions per recording session were quantified to assess data yield. In the standard 8–32 kHz dataset, we identified distinct populations of sound-activated and sound-suppressed neurons summed from all sessions (Figure 4A). As shown in the grand average traces, the activated populations exhibited a sharp, time-locked onset at *t* = 0, confirming precise synchronization between behavioral events and imaging frames. Furthermore, the cell responsiveness distributions indicate a consistent yield of high-quality neurons in every recording session. We observed similarly high-quality dynamics in the shifted frequency conditions of 4–16 kHz and 7–28 kHz (Figure 4B, C). Across all datasets, the neuronal populations maintained high signal-to-noise ratios with peak *Z*-scores *>* 1.5 and displayed the expected biological heterogeneity in the proportions of activated versus suppressed cells[18]. Collectively, these results confirm that the recordings capture robust, biologically relevant neural dynamics across diverse experimental contexts.

**Figure 4:**
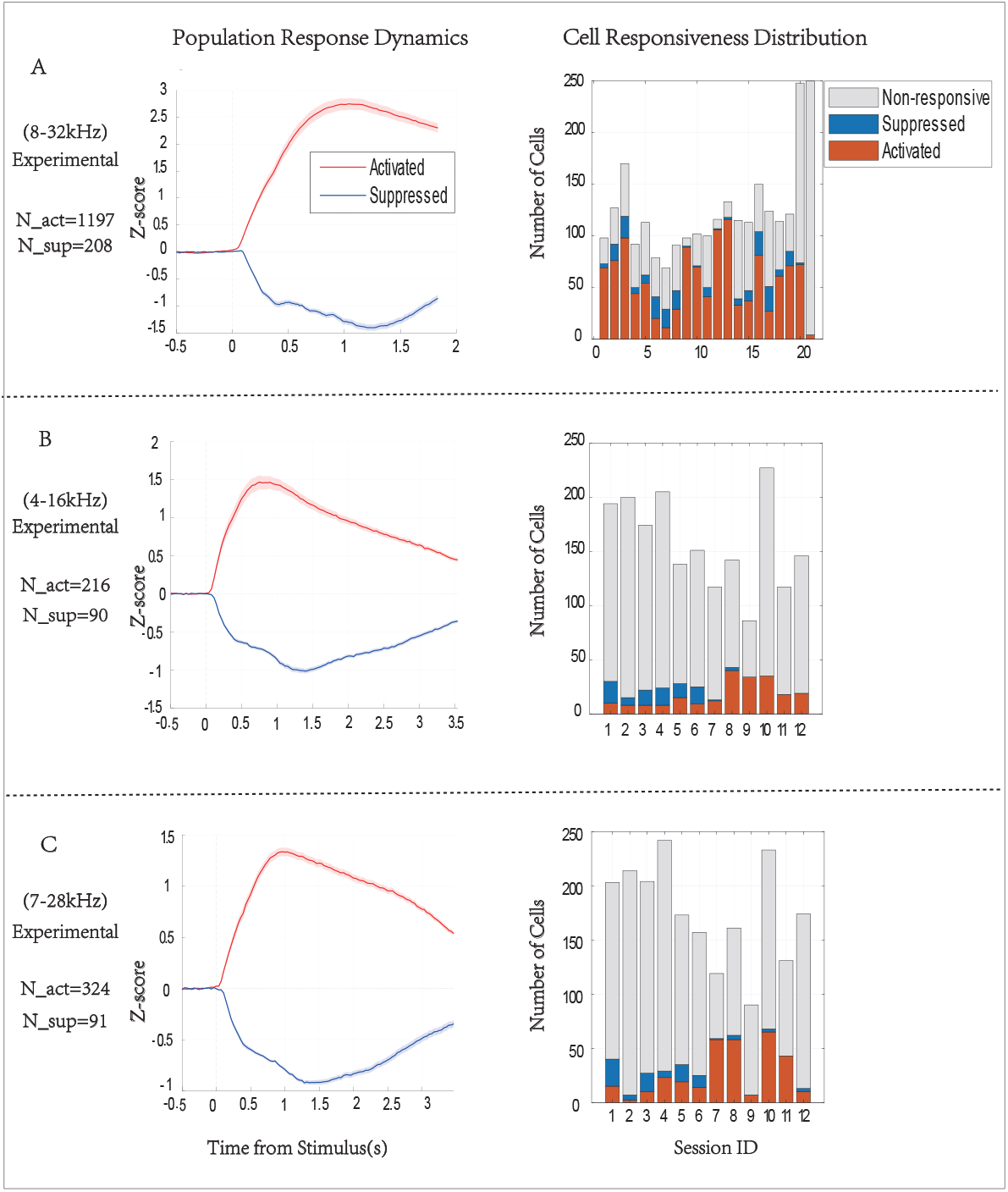
Overview of neuronal responsiveness and population dynamics across three experimental groups. Analysis of sound-evoked activity. Left column shows the grand average temporal dynamics for sound-activated and sound-suppressed neurons. The vertical dashed line at *t* = 0 denotes stimulus onset. Total cell counts are annotated below each panel. Right column illustrates the session-wise distribution of cell responsiveness. Stacked bars indicate the number of activated, suppressed, and non-responsive neurons per recording session.

#### Verification of Spectral Tuning

Beyond general responsiveness, a critical quality metric for auditory cortex datasets is the preservation of frequency selectivity [19]. We examined the tuning properties of the core population of sound-activated neurons identified across all three experimental groups (Figure 5).

**Figure 5:**
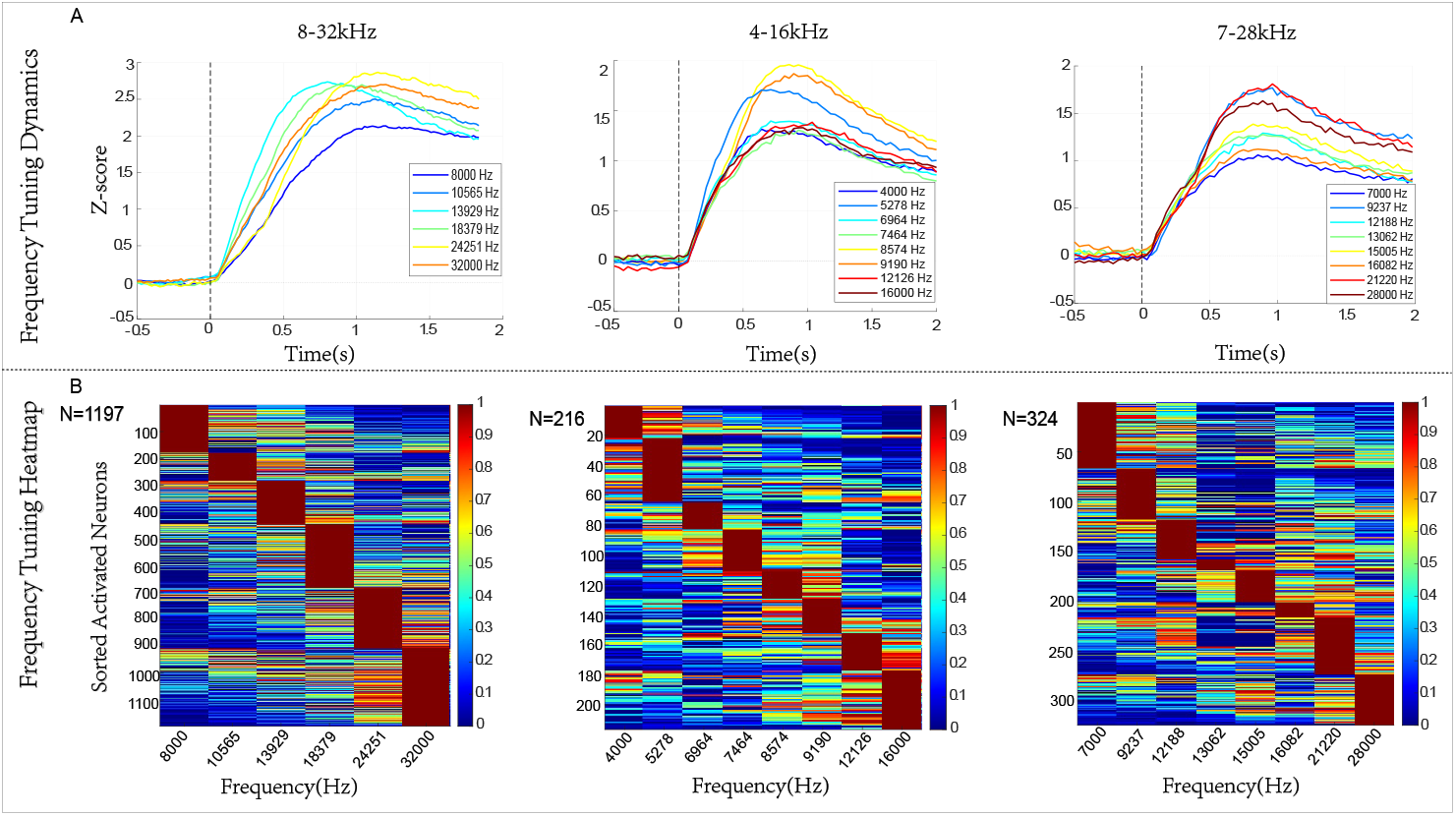
Frequency selectivity and tonotopic organization of sound-activated neurons. Traces on the top panels show the mean *Z*-scored activity of activated neurons for each presented tone frequency across three experimental conditions. The vertical dashed line denotes stimulus onset (*t* = 0). Heatmaps in the bottom demonstrate population frequency tuning. Each chart represents the normalized tuning curve of a single activated neuron, sorted by its best frequency [21]. Total neuron counts are annotated in each panel.

We first analyzed the temporal profiles of the population response grouped by stimulus frequency (Figure 5A). To generate these dynamics, the *Z*-scored activity of all activated neurons was simply averaged across both cells and trials for each sound frequency. In all three datasets, the population activity was strongly modulated by sound frequency. Specifically, in the standard band of 8–32 kHz and low-shift condition of 4–16 kHz, intermediate frequencies elicited the strongest population responses. Interestingly, the high-shift condition of 7–28 kHz exhibited a distinct bimodal response profile, with the most robust population activity driven by two non-adjacent frequencies: a lower-mid tone (9237 Hz) and a high tone (21220 Hz). This bimodal distribution at the pooled population level likely arises from the anatomical heterogeneity of the sampled fields of view. Specifically, the two-photon imaging windows in this particular dataset may have been preferentially centered over both the low frequency and high frequency regions of A1, leading to an overrepresentation of these specific frequency bands in the grand average [20].

To visualize individual neuron selectivity, we constructed frequency tuning heatmaps. For each activated neuron, we calculated its mean *Z*-scored response to each frequency during the 1.5 s post-stimulus window. These individual tuning curves were then min-max normalized and sorted by their Best Frequency (BF), defined as the specific sound frequency eliciting the maximal neural response magnitude[21]. As shown in Figure 5B, a striking diagonal structure emerges across all datasets. This indicates that the vast majority of activated neurons have a unique and distinct best frequency, rather than responding broadly to all tones. This distinct clustering confirms that the dataset successfully captures the fine-grained tonotopic organization of the primary auditory cortex, making it highly suitable for decoding analyses and stimulus reconstruction tasks.

### Dataset Evaluation

To evaluate the potential of the prefrontal cortex calcium imaging dataset constructed in this study for supporting neural decoding research, we conducted a standardized benchmarking test. The test aimed to quantitatively assess the performance of machine learning models in decoding two key behavioral variables from the standardized neural signals of this dataset: discriminating the auditory stimulus type and predicting the behavioral choice. Specifically, we systematically compared four models: Multilayer Perceptron (MLP), Support Vector Machine (SVM), Convolutional Neural Network (CNN), and Long Short-Term Memory network (LSTM), to examine the dataset’s compatibility with different computational methods.

The raw data underwent standardized preprocessing to ensure fairness and reproducibility of the evaluation. First, to address differences in original acquisition frame rates (28 Hz or 55 Hz), all neural time series were extracted from an identical 1-second post-stimulus time window and uniformly resampled to 55 Hz via linear interpolation. Next, we unified the varying number of recorded neurons across sessions by a cropping strategy. It was employed to adjust the number of regions of interest for all samples to a global minimum. After preprocessing, the data were split into a training set and an independent test set, with 10-fold cross-validation applied to the training set for hyperparameter optimization, and the final model performance was evaluated on the test set.

The models employed in the evaluation and their core configurations were as follows. The model of MLP comprised three fully-connected hidden layers with 256, 128, and 64 neurons, utilizing the LeakyReLU activation function and dropout rate of 0.3. In SVM, we leveraged radial basis function kernel with the regularization parameter *C* = 1.0. As for CNN, the neural activity data is shaped into single-channel 2D images for processing with 2D convolutional layers. LSTM was set as a four-layer unidirectional network. All deep learning models were trained using the Adam optimizer with learning rate of 0.001 with an early stopping mechanism. The evaluation used the test set classification accuracy as the primary metric.

The model performance comparison is shown in Figure 6. MLP and SVM achieved performance that was significantly superior to that of both CNN and LSTM on both neural decoding tasks. Specifically, the test accuracies of discriminating the auditory stimulus type for MLP and SVM were 77.45% and 75.75%, respectively; for the task of predicting action choices, the accuracies were 78.30% and 76.23%, respectively.

**Figure 6:**
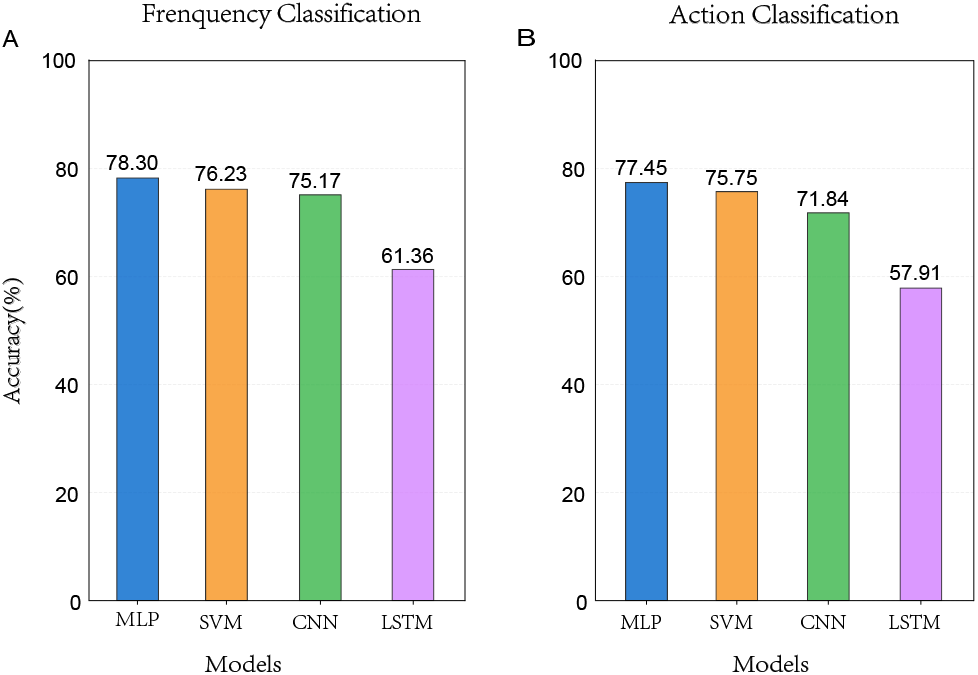
Performance comparison of MLP, SVM, CNN, and LSTM models on two decoding tasks, discrimination of stimulus types and action choices. Bar heights represent classification accuracy on independent test sets.

## Usage Notes

The dataset presented in our study offers a unique resource for bridging biological intelligence and artificial intelligence. By providing high-dimensional, trial-aligned neural dynamics paired with precise behavioral outputs, these data are particularly suitable for two emerging research directions: neural decoding and brain activity simulation.

### Neural Decoding

The simultaneous recording of large-scale population activity in A1 (up to 150 neurons per session) during a decision-making task makes this dataset ideal for developing and benchmarking neural decoders. Researchers can utilize the traces for neural decoding. On the one hand, it can be used to train classifiers to reconstruct the auditory stimulus frequency from single-trial population vectors. On the other hand, researchers can apply deep learning models to predict the animal’s learnable choice based on the temporal evolution of neural activity. Recent studies have demonstrated that such decoding models can reveal the internal dynamics of decision-making [22, 19, 23]. Moreover, since the dataset includes error trials and miss trials, researchers can investigate the neural correlates of lapses in attention or decision errors, a critical step in understanding the variability of biological computation [15].

### Data-Driven Brain Simulation

Beyond decoding, this dataset serves as ground truth for training data-driven dynamical systems models. In the field of latent dynamics modeling, tools like LFADS (Latent Factor Analysis via Dynamical Systems) [24] or CEBRA [25] can be applied to these data to extract low-dimensional latent trajectories that govern the population activity. The dataset can also be used to train biologically constrained models to simulate the same auditory discrimination task. By comparing the hidden activity of the artificial network with the recorded A1 activity, researchers can test hypotheses about the computational algorithms implemented by cortical circuits [26, 27].

## Data Availability

The raw data can be found on the website of the brain data center (https://www.braindatacenter.cn/datacenter/web/#/dataSet/details?id=1722080685634592769). The data samples and examples of utilizing samples to implement decoding are specified on a GitHub page (https://github.com/YuYongZhen98/Calcium-Imaging-in-Decision-Making-Mice).

## Code availability

The code for importing and processing the data described in this paper, as well as the code for generating the figures and example processing, is available in the public repository: https://github.com/YuYongZhen98/Calcium-Imaging-in-Decision-Making-Mice.

## Funding

This work was supported by the Brain Science and Brain-like Intelligence Technology - National Science and Technology Major Project (2025ZD0217200), Strategic Priority Research Program of the Chinese Academy of Sciences (Grant No. XDB1010302), CAS Project for Young Scientists in Basic Research (YSBR-116), Youth Innovation Promotion Association CAS, Shanghai Leading Talent Program of Eastern Talent Plan, the Lingang Laboratory Fund (Grant No. LG-GG-202402-06-07, LGL-1987-09), the Shanghai Municipal Science and Technology Project (Grant No. 25ZR1401370, 25LN3200400), Special Support Project of Guangdong Province (Grant No.0720240209). The numerical calculations in this study were carried out on the ORISE Supercomputer.

## Competing Interests

The authors declare no competing interests.

## Author Contributions

B.H. and L.L. co-designed the experiments and developed the code for data processing. Y.Y. developed the machine learning models discussed in the section of Modeling Performance and co-designed the experiments. L.L. assisted with the data analysis and contributed to the statistical analysis of the dataset. Y.X. and Y.Z. acquired the data. N.X. supervised the data collection. T.Z. supervised the overall project, contributed to the study design, and provided critical guidance throughout the experimental and writing phases. All authors contributed to the conception of the study, provided critical feedback on the experimental design and data analysis, and participated in the writing and review of the manuscript.

## Notes

### Competing Interest Statement

The authors have declared no competing interest.

## References

[1] Winfried Denk, James H Strickler, and Watt W Webb. “Two-photon laser scanning fluorescence microscopy”. In: Science 248.4951 (1990), pp. 73–76.

[2] Christine Grienberger and Arthur Konnerth. “Imaging calcium in neurons”. In: Neuron 73.5 (2012), pp. 862–885.

[3] Tsai-Wen Chen et al. “Ultrasensitive fluorescent proteins for imaging neuronal activity”. In: Nature 499.7458 (2013), pp. 295–300.

[4] Nicholas James Sofroniew et al. “A large field of view two-photon mesoscope with subcellular resolution for in vivo imaging”. In: elife 5 (2016), e14472.

[5] Petr Znamenskiy and Anthony M Zador. “Corticostriatal neurons in auditory cortex drive decisions during auditory discrimination”. In: Nature 497.7450 (2013), pp. 482–485.

[6] Jonathan B Fritz et al. “Auditory attention—focusing the searchlight on sound”. In: Current opinion in neurobiology 17.4 (2007), pp. 437–455.

[7] Stephen V David, Jonathan B Fritz, and Shihab A Shamma. “Task reward structure shapes rapid receptive field plasticity in auditory cortex”. In: Proceedings of the National Academy of Sciences 109.6 (2012), pp. 2144–2149.

[8] Simon Musall et al. “Single-trial neural dynamics are dominated by richly varied movements”. In: Nature neuroscience 22.10 (2019), pp. 1677–1686.

[9] Santiago Jaramillo and Anthony Zador. “Auditory cortex mediates the perceptual effects of acoustic temporal expectation”. In: Nature Precedings (2010), pp. 1–1.

[10] Jeremy G Turner et al. “Hearing in laboratory animals: strain differences and nonauditory effects of noise”. In: Comparative medicine 55.1 (2005), p. 12.

[11] Santiago Jaramillo and Anthony M Zador. “Mice and rats achieve similar levels of performance in an adaptive decision-making task”. In: Frontiers in systems neuroscience 8 (2014), p. 173.

[12] Anthony Zador et al. “Toward next-generation artificial intelligence: Catalyzing the neuroai revolution”. In: arXiv preprint arXiv:2210.08340 (2022).

[13] Joshua I Glaser et al. “Machine learning for neural decoding”. In: eneuro 7.4 (2020).

[14] Jesse A Livezey and Joshua I Glaser. “Deep learning approaches for neural decoding across architectures and recording modalities”. In: Briefings in bioinformatics 22.2 (2021), pp. 1577–1591.

[15] Yu Xin et al. “Sensory-to-category transformation via dynamic reorganization of ensemble structures in mouse auditory cortex”. In: Neuron 103.5 (2019), pp. 909–921.

[16] Zengcai V Guo et al. “Procedures for behavioral experiments in head-fixed mice”. In: PloS one 9.2 (2014), e88678.

[17] Naoshige Uchida and Zachary F Mainen. “Speed and accuracy of olfactory discrimination in the rat”. In: Nature neuroscience 6.11 (2003), pp. 1224–1229.

[18] Hiroyuki K Kato, Shea N Gillet, and Jeffry S Isaacson. “Flexible sensory representations in auditory cortex driven by behavioral relevance”. In: Neuron 88.5 (2015), pp. 1027–1039.

[19] Carsen Stringer et al. “Spontaneous behaviors drive multidimensional, brainwide activity”. In: Science 364.6437 (2019), eaav7893.

[20] Gideon Rothschild, Israel Nelken, and Adi Mizrahi. “Functional organization and population dynamics in the mouse primary auditory cortex”. In: Nature neuroscience 13.3 (2010), pp. 353–360.

[21] Gideon Rothschild, Israel Nelken, and Adi Mizrahi. “Functional organization and population dynamics in the mouse primary auditory cortex”. In: Nature neuroscience 13.3 (2010), pp. 353–360.

[22] Felix Pei et al. “Neural latents benchmark’21: evaluating latent variable models of neural population activity”. In: arXiv preprint arXiv:2109.04463 (2021).

[23] Binjie Hong et al. “Bidirectional cross-day alignment of neural spikes and behavior using a hybrid SNN-ANN algorithm”. In: Frontiers in Neuroscience 20 (2026), p. 1772958.

[24] Chethan Pandarinath et al. “Inferring single-trial neural population dynamics using sequential auto-encoders”. In: Nature methods 15.10 (2018), pp. 805–815.

[25] Steffen Schneider, Jin Hwa Lee, and Mackenzie Weygandt Mathis. “Learnable latent embeddings for joint behavioural and neural analysis”. In: Nature 617.7960 (2023), pp. 360–368.

[26] Guangyu Robert Yang and Xiao-Jing Wang. “Artificial neural networks for neuroscientists: a primer”. In: Neuron 107.6 (2020), pp. 1048–1070.

[27] Saurabh Vyas et al. “Computation through neural population dynamics”. In: Annual review of neuroscience 43.1 (2020), pp. 249–275.

